# MKado: a toolkit for McDonald-Kreitman tests of natural selection

**DOI:** 10.64898/2026.03.02.709122

**Authors:** Angel G. Rivera-Colón, Clara T. Rehmann, Andrew D. Kern

## Abstract

**Summary:** `MKado` is a Python toolkit for performing McDonald-Kreitman (MK) tests of natural selection from aligned coding sequences. It implements the standard MK test as well as a wide variety of its extensions and related statistics, including a number of methods for estimating the fraction of adaptive substitutions (*α*) while accounting for slightly deleterious mutations, with a unified command-line interface and Python API. `MKado` supports parallel batch processing of thousands of genes with near-linear scaling, and provides publication-ready visualizations including volcano plots and asymptotic *α* curves.

**Availability and Implementation:** `MKado` is freely available at https://github.com/kr-colab/mkado under the MIT license. Full documentation is available at https://mkado.readthedocs.io. `MKado` is implemented in Python and installable via `pip`.

## Introduction

The McDonald-Kreitman (MK) test (McDonald and Kreitman, 1991) is among the most widely used methods for detecting natural selection at the molecular level. By comparing the ratio of nonsynonymous to synonymous changes within species (polymorphism) to that between species (divergence), the MK test provides a direct test of the neutral theory prediction that these ratios should be equal. Departures from this expectation, often quantified by the neutrality index (NI; Rand and Kann, 1996) or the fraction of adaptive substitutions *α* (Smith and Eyre-Walker, 2002), have revealed pervasive positive selection across a wide range of taxa (e.g., Lozada-Chávez et al., 2025; Brown et al., 2024; Liti et al., 2009; Lam et al., 2010; Downing et al., 2009; Hughes et al., 2008; Li et al., 2008).

A key challenge for the standard MK test is that slightly deleterious nonsynonymous mutations, which contribute to polymorphism but rarely fix, inflate the number of nonsynonymous polymorphisms relative to nonsynonymous substitutions and bias *α* downward. Early approaches addressed this by excluding low-frequency polymorphisms, either by distinguishing tip from interior mutations on the gene tree (Templeton, 1996) or by applying a derived allele frequency cutoff (Fay et al., 2001). The asymptotic MK test further addresses this limitation by estimating *α* as a function of derived allele frequency and extrapolating to fixation, effectively removing the contribution of weakly deleterious variants (Messer and Petrov, 2013; Haller and Messer, 2017). Rather than discarding low-frequency polymorphisms entirely, the imputed MK test uses the ratio of low-to high-frequency synonymous polymorphisms to estimate the expected number of weakly deleterious nonsynonymous polymorphisms and subtracts them from the total, retaining more data while correcting for the bias (Murga-Moreno et al., 2022). To address lineage-specific selection, the polarized MK test uses a second outgroup to assign fixed differences to specific lineages, allowing lineage-specific inference of selection (Begun et al., 2007).

Despite the centrality of MK-based methods to molecular evolution, existing software implementations remain fragmented. `DnaSP` (Librado and Rozas, 2009) provides the standard test through a Windows-only graphical interface. The `iMKT` R package and web server (Murga-Moreno et al., 2019) implements several extensions but requires pre-computed summary statistics and is tightly coupled to specific databases. The `asymptoticMK` web tool (Haller and Messer, 2017) implements the asymptotic test but does not accept raw sequence data. The `degenotate` software, part of the `snpArcher` pipeline Mirchandani et al. (2023), can calculate both standard and imputed versions of the test; however it does so only using genetic variants in VCF format and is thus dependent on (and susceptible to) the liftover of outgroup sequences. No existing tool provides a unified command-line interface that takes aligned coding sequences as input and supports the standard, polarized, and asymptotic MK tests together with batch processing across thousands of genes (Table 1).

**Table 1:**
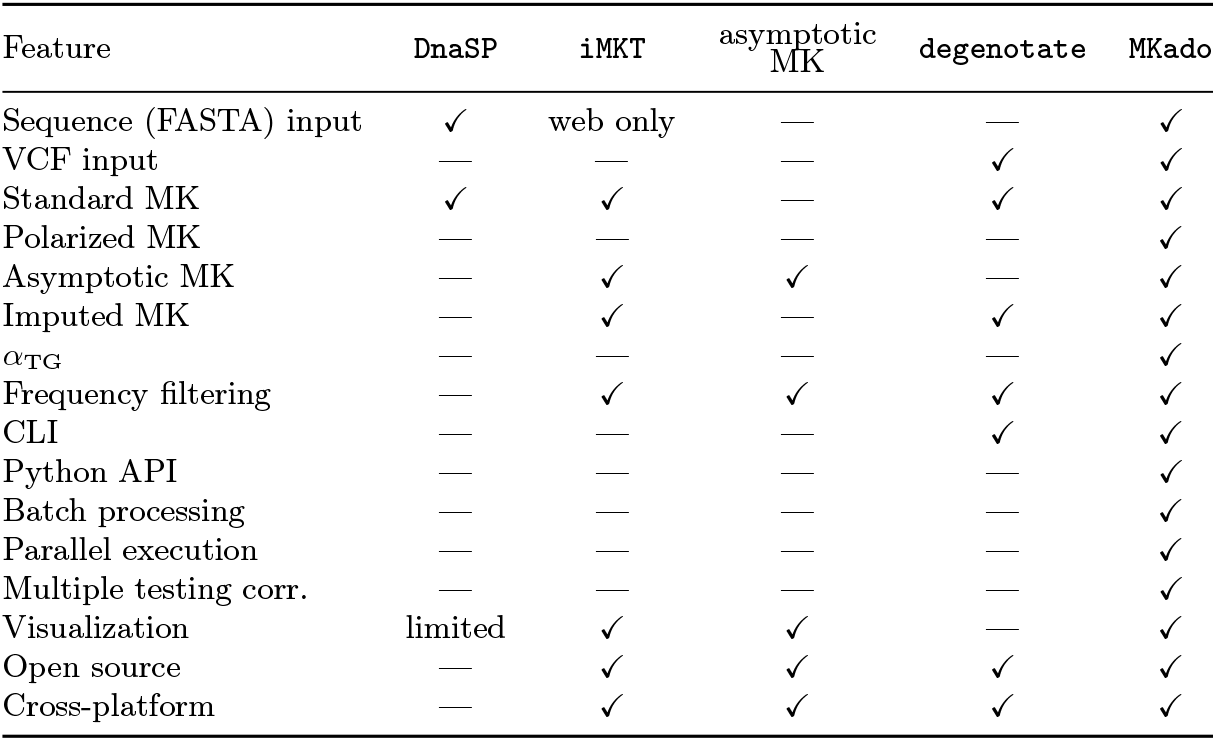
Feature comparison of MK test software. Checkmarks indicate built-in support; dashes indicate the feature is absent.

| Feature | DnaSP | iMKT | asymptotic<br>MK | degenotate | MKado |
| --- | --- | --- | --- | --- | --- |
| Sequence (FASTA) input | ✓ | web only | — | — | ✓ |
| VCF input | — | — | — | ✓ | ✓ |
| Standard MK | ✓ | ✓ | — | ✓ | ✓ |
| Polarized MK | — | — | — | — | ✓ |
| Asymptotic MK | — | ✓ | ✓ | — | ✓ |
| Imputed MK | — | ✓ | — | ✓ | ✓ |
| $\alpha_{TG}$ | — | — | — | — | ✓ |
| Frequency filtering | — | ✓ | ✓ | ✓ | ✓ |
| CLI | — | — | — | ✓ | ✓ |
| Python API | — | — | — | — | ✓ |
| Batch processing | — | — | — | — | ✓ |
| Parallel execution | — | — | — | — | ✓ |
| Multiple testing corr. | — | — | — | — | ✓ |
| Visualization | limited | ✓ | ✓ | — | ✓ |
| Open source | — | ✓ | ✓ | ✓ | ✓ |
| Cross-platform | — | ✓ | ✓ | ✓ | ✓ |

Here we present `MKado`, a Python package that fills this gap. `MKado` takes both codon-aligned FASTA and VCF files as input and implements the standard, polarized, asymptotic, and imputed MK tests, as well as related statistics, with a consistent command-line interface. It supports parallel batch processing of genome-scale datasets, automatic multiple testing correction, and built-in visualization including volcano plots and asymptotic *α* curves. `MKado` is installable via `pip` and is designed to integrate naturally with modern Python-based bioinformatics workflows.

### Features and Implementation

`MKado` is implemented in Python and provides both a command-line interface built on `Typer` and a Python API for programmatic use. It accepts codon-aligned FASTA files as input, requiring no pre-processing or external format conversion. Sequences are automatically partitioned into ingroup and outgroup sets by name pattern matching, supporting both multi-species alignments in a single FASTA file and separate files containing ingroup or outgroup sequences. `MKado` also accepts variant-call input via the `vcf` subcommand, which uses a multi-sample ingroup VCF, a single-sample outgroup VCF, a reference FASTA, and a GFF3 annotation to derive codon-level polymorphism and divergence counts directly and supports the full set of MK analyses available to the FASTA workflow. Together, these two input modes distinguish `MKado` from FASTA-only tools (`DnaSP`, `asymptoticMK`) and from VCF-only tools (`degenotate`, `fastDFE`). Full documentation, including tutorials and API reference, is available at https://mkado.readthedocs.io.

#### Analysis methods

`MKado` implements five MK-based analyses. The *standard* test computes the classical 2 × 2 table of nonsynonymous and synonymous changes in polymorphism and divergence, with significance assessed by Fisher’s exact test or the *G*-test. *D*_*n*_ and *D*_*s*_ are the numbers of fixed differences between ingroup and outgroup at codon positions where the inferred substitution is nonsynonymous or synonymous, respectively, under the supplied genetic code. *P*_*n*_ and *P*_*s*_ are the corresponding numbers of ingroup polymorphic sites at which the derived (non-outgroup) allele segregates above an optional frequency threshold (default zero, i.e. all polymorphisms; `--min-freq` *X* excludes sites with derived allele frequency below *X*, and `--no-singletons` excludes singletons by setting the threshold to 1*/n* for sample size *n*). By default *P*_*n*_ and *P*_*s*_ follow the `DnaSP` convention and exclude sites that also segregate in the outgroup; `--pool-polymorphisms` pools ingroup and outgroup polymorphism instead. Summary statistics include the neutrality index (NI; Rand and Kann, 1996), *α* (Smith and Eyre-Walker, 2002), and the Direction of Selection statistic (DoS; Stoletzki and Eyre-Walker, 2011). Tests that depend on derived allele frequency—the asymptotic and imputed MK tests described below—require knowing the derived allele at each polymorphic site. `MKado` polarizes alleles automatically using the outgroup: at each polymorphic site, the allele matching the outgroup is treated as ancestral and any non-outgroup allele as derived. This automatic single-outgroup polarization is distinct from the *polarized* MK test below, which uses a *second* outgroup to attribute each fixed difference to either the ingroup or the outgroup lineage. The *polarized* test uses a second outgroup to infer the ancestral state at each fixed difference, assigning divergence to the ingroup or outgroup lineage for lineage-specific inference. The *asymptotic* test bins polymorphisms by derived allele frequency, computes *α* within each bin, and fits an exponential model *α*(*x*) = *a* + *be*^−*cx*^ to extrapolate the asymptotic *α* at *x* = 1, correcting for the contribution of weakly deleterious variants (Messer and Petrov, 2013). A linear model is also fitted and AIC selects the better specification, following Haller and Messer (2017). `MKado` supports both the original per-bin SFS construction of Messer and Petrov (2013), in which *P*_*n*_(*x*) and *P*_*s*_(*x*) count polymorphic sites with derived allele frequency *equal to x*, and the cumulative inclusive variant of Uricchio et al. (2019), in which *P*_*n*_(*x*) and *P*_*s*_(*x*) count polymorphic sites with derived allele frequency *at or above x* (selectable with `--sfs-mode {at,above}`); the cumulative form has the same asymptote at *x* = 1 but is more robust to bin-count sparsity in large samples. The asymptotic test supports both per-gene and aggregated (pooled across genes) modes; for the aggregated mode, two confidence-interval methods are available (`--ci-method`): a parametric Monte-Carlo procedure that samples the curve-fit covariance matrix (default; fast) and a case-resampling bootstrap that resamples the pooled polymorphism list and refits per replicate (slower per replicate but more principled when per-bin counts are small). The *imputed* test corrects for weakly deleterious mutations by estimating their contribution from the ratio of low-to high-frequency synonymous polymorphisms (Murga-Moreno et al., 2022). Polymorphisms are split at a derived allele frequency cutoff (default 15%), and the expected number of weakly deleterious nonsynonymous polymorphisms below the cutoff is imputed as *P*_*wd*_ = *P*_*n*,low_ − *P*_*n*,high_ × (*P*_*s*,low_*/P*_*s*,high_). The corrected count 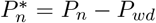 is then used in the standard 2 × 2 test to obtain a bias-corrected *α*. This approach retains more data than simple frequency filtering while still removing the downward bias from slightly deleterious variants. Finally, the Tarone-Greenland *α*_TG_ estimator (Stoletzki and Eyre-Walker, 2011) provides an unbiased multi-gene estimate of *α* by weighting each gene inversely by *P*_*s*_ + *D*_*s*_, correcting for sample size heterogeneity across loci.

#### Codon change classification

When comparing codons that differ at multiple positions, the assignment of changes as synonymous or nonsynonymous is not straightforward because the order of substitutions is unknown. `MKado` pre-computes all shortest mutational paths between every pair of codons and weights the synonymous and nonsynonymous contributions equally across paths, following the approach of Nei and Gojobori (1986).

#### Batch processing and output

MKado’s `batch` command processes directories of alignment files in parallel using Python’s multiprocessing facilities with configurable worker counts. Benjamini-Hochberg false discovery rate (FDR) *q*-values are computed automatically across all genes. Results are available in human-readable, TSV, or JSON formats. A companion `info` command reports alignment diagnostics (sequence count, length, and codon structure) for input validation.

To evaluate scaling performance, we benchmarked `MKado` on 12,437 human protein-coding genes aligned with chimpanzee and orangutan orthologs from `OrthoMaM` (Allio et al., 2024), with polymorphism data from 178 Yoruba individuals in the 1000 Genomes Project (1000 Genomes Project Consortium, 2015). Processing all genes with 128 worker threads completed in 28 seconds, a ∼50× speedup over single-threaded execution. Per-gene work (alignment parsing, codon classification, and the per-gene MK computation) takes only milliseconds, so the per-process spawn overhead from Python’s `multiprocessing` grows non-negligible relative to per-task work as the worker count rises. Figure S1 shows approximately linear scaling up to roughly 16 workers, with progressively diminishing efficiency past 32 workers; we therefore recommend setting the worker count well below the gene count for batch runs at any scale. For workloads dominated by aggregated tests (asymptotic MK with case-resampling bootstrap CIs), where per-task work is much larger, scaling extends usefully to higher worker counts (Fig. S3).

#### Visualization

`MKado` generates publication-ready plots directly from the command line. Volcano plots display − log_10_(NI) against − log_10_(*p*) for each gene, with Bonferroni and nominal significance thresholds (left panel of Fig. 1). The asymptotic test produces *α*(*x*) curves showing the fitted model and bootstrap confidence intervals (right panel of Fig. 1).

**Figure 1.**
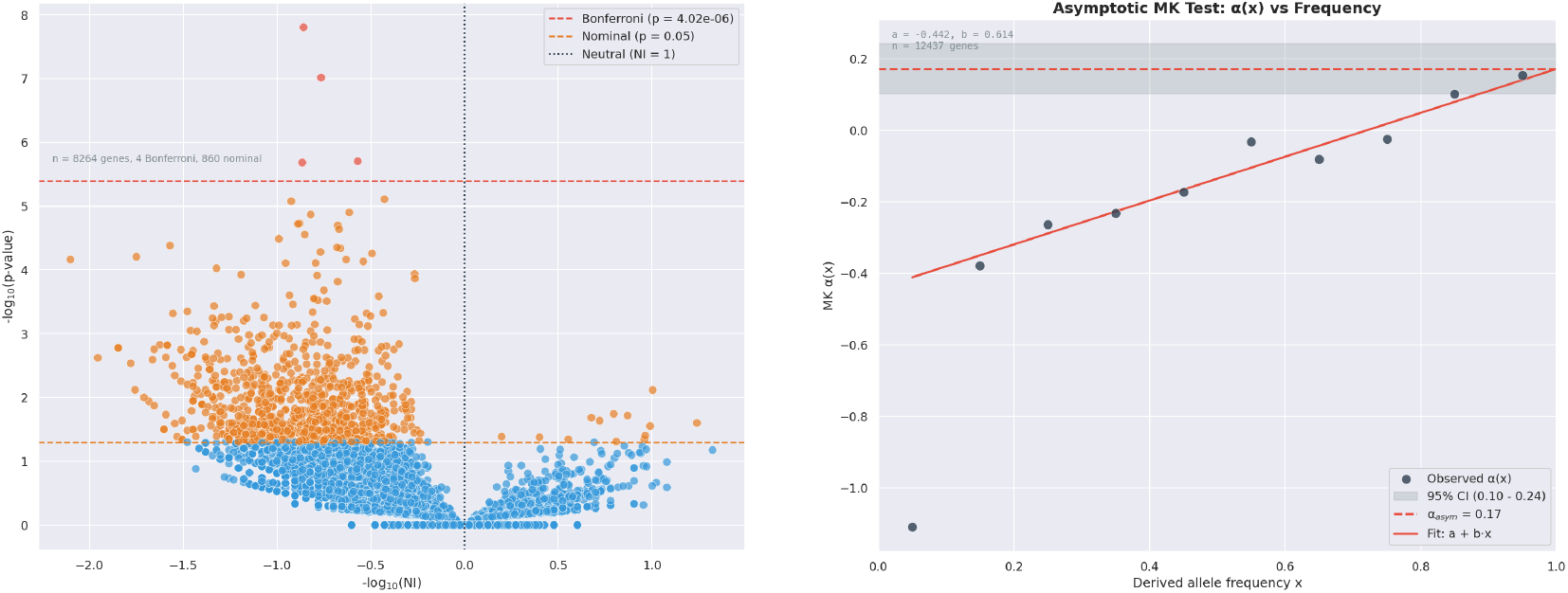
MK test results for 12,437 human protein-coding genes using *Pongo abelii* as the outgroup for divergence. Left: Volcano plot from per-gene standard MK tests. Red points are significant after Bonferroni correction, while orange points are nominally significant (*p <* 0.05). The dashed lines indicate Bonferroni and nominal thresholds. The leftward shift of the cloud (NI *>* 1) reflects the wellknown excess of weakly deleterious nonsynonymous polymorphism in humans. Right: Asymptotic *α* curve for pooled data. Points represent the observed *α* at each derived allele frequency cutoff; the solid line represents the fitted curve, in this case the linear model selected by AIC. The shaded area indicates the 95% confidence interval around the fitted asymptotic *α* value.

### Application

To demonstrate `MKado` at genome scales, we assembled a dataset of 12,437 human protein-coding genes with both polymorphism and divergence data. Codon-aligned orthologous sequences for human (*Homo sapiens*), chimpanzee (*Pan troglodytes*), and orangutan (*Pongo abelii*) were obtained from `OrthoMaM` v12 (Allio et al., 2024), which provides one canonical coding-sequence ortholog per gene across mammalian taxa. Human intraspecific polymorphism was drawn from 178 Yoruba (YRI) individuals in the 1000 Genomes Project NYGC high-coverage dataset (1000 Genomes Project Consortium, 2015), yielding 356 phased haplotypes per gene.

For each gene, polymorphism was projected onto the `OrthoMaM` alignment in five steps: (i) the canonical Ensembl CDS for the gene was reconstructed from the GRCh38 reference and the GTF-derived exon set, providing the mapping from CDS position to genomic coordinate; (ii) the ungapped human row of the `OrthoMaM` alignment was matched against the Ensembl CDS to obtain the alignment-column to CDS-position correspondence (genes for which this match failed, typically because of curation differences between `OrthoMaM` and Ensembl, were dropped); (iii) for every CDS position that survived this match, the overlapping YRI variants were extracted from the 1000 Genomes NYGC VCF, with reference-allele identity confirmed and minus-strand alleles complemented to match the CDS orientation; (iv) per-individual phased haplotypes were assembled from the extracted variants, yielding 356 human haplotype sequences over the Ensembl CDS positions retained by `OrthoMaM`; and (v) these haplotypes were re-gapped to the `OrthoMaM` alignment, dropping any human-CDS columns absent from the `OrthoMaM` alignment and inserting the alignment’s existing gap structure (so insertions, deletions, and non-aligned regions in the multi-species alignment are preserved). Chimpanzee and orangutan sequences were taken directly from the `OrthoMaM` alignment. Multi-transcript genes are reduced to the single canonical CDS chosen by the `OrthoMaM` curation; alternative transcripts are not analysed separately. The resulting per-gene multi-species FASTAs are codon-aligned and ready for direct input to `MKado`.

Using orangutan as the outgroup, genome-wide pooled counts were *D*_*n*_ = 94,578, *D*_*s*_ = 188,325, *P*_*n*_ = 44,850, and *P*_*s*_ = 47,838. The per-gene standard MK test identified numerous genes with significant departures from neutrality, including genes significant after Bonferroni correction (volcano plot, left panel of Fig. 1). As expected, the majority of testable genes showed NI *>* 1, reflecting the well-characterized excess of slightly deleterious nonsynonymous polymorphism in humans (Li et al., 2008). Using chimpanzee as the outgroup yielded fewer fixed differences (*D*_*n*_ = 37,120, *D*_*s*_ = 65,228), roughly 2.5-fold less than orangutan (left panel of Fig. S2), and no individual genes survived Bonferroni correction, consistent with the limited power of single-gene tests at shorter divergence times.

The Tarone-Greenland weighted estimate (Stoletzki and Eyre-Walker, 2011) across all genes was strongly negative for both outgroups: 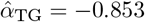 (95% CI: −0.888 to −0.812) for orangutan and 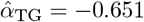 (95% CI: −0.682 to −0.607) for chimpanzee, with corresponding NI_TG_ values of 1.853 and 1.651. These negative values confirm the pervasive excess of weakly deleterious nonsynonymous polymorphism in humans. Excluding singletons shifted both estimates upward (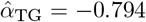 for orangutan; −0.606 for chimpanzee), as expected given the enrichment of slightly deleterious variants at low frequency. Applying a 15% derived allele frequency cutoff (Fay et al., 2001) brought both estimates close to zero (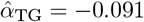 for orangutan; −0.041 for chimpanzee), removing the bulk of low-frequency deleterious variants but still failing to recover a signal of positive selection. Similarly, the aggregated imputed MK test (Murga-Moreno et al., 2022), using a 15% derived allele frequency cutoff, estimated 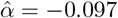 for orangutan and 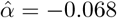 for chimpanzee, substantially reducing the downward bias but likewise not recovering a positive signal of adaptation.

In contrast, the aggregated asymptotic MK test, which models the frequency dependence of *α* and extrapolates to fixation (Messer and Petrov, 2013), successfully recovered a positive signal of adaptive evolution, estimating 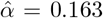 (95% CI: 0.075 to 0.253; right panel of Fig. 1) using orangutan as the outgroup, indicating that approximately 16% of amino acid substitutions on the human lineage were driven by positive selection. With chimpanzee as the outgroup the estimate widened to 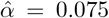 (95% CI: −1.22 to 0.11; right panel of Fig. S2), illustrating the greater uncertainty from the shallower divergence, but also indicating a potential biological signal of less frequent adaptive evolution after the human-Pongo split. These results are consistent with previous analyses of human coding sequences (Messer and Petrov, 2013).

The entire analysis—standard MK tests, Tarone-Greenland estimation, asymptotic estimation, imputed estimation, and volcano plot generation for all 12,437 genes—completed in under one minute on a multi-core workstation using `MKado`’s parallel batch mode.

## Acknowledgements

We thank Matt Hahn, Dan Schrider, and Scott Small for comments on an earlier version of this manuscript and various members of the Kern-Ralph colab for helpful discussions and feedback.

