## Supplemental Figures for "MKado: a toolkit for McDonald-Kreitman tests of natural selection"

### Supplementary Results

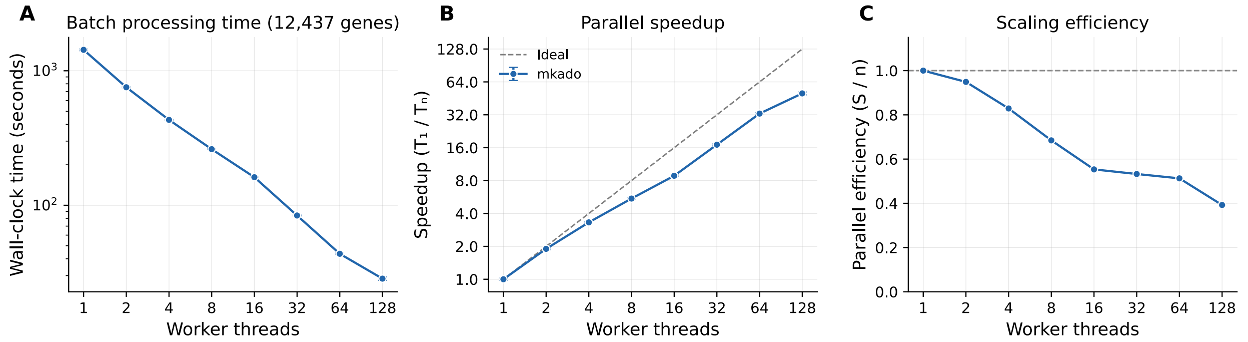


Figure S1: Parallel scaling of `MKado` batch processing on 12,437 human protein-coding genes. (**A**) Wall-clock time versus worker threads. (**B**) Speedup relative to single-threaded execution, with ideal linear scaling shown as a dashed line. (**C**) Parallel efficiency (speedup divided by thread count). Real scaling diverges from the ideal linear curve because the per-gene work (parsing, codon classification, and the per-gene MK test) takes on the order of milliseconds, while the spawn cost of a Python `multiprocessing` worker is fixed; the spawn overhead therefore grows non-negligible relative to per-task work at high worker counts.

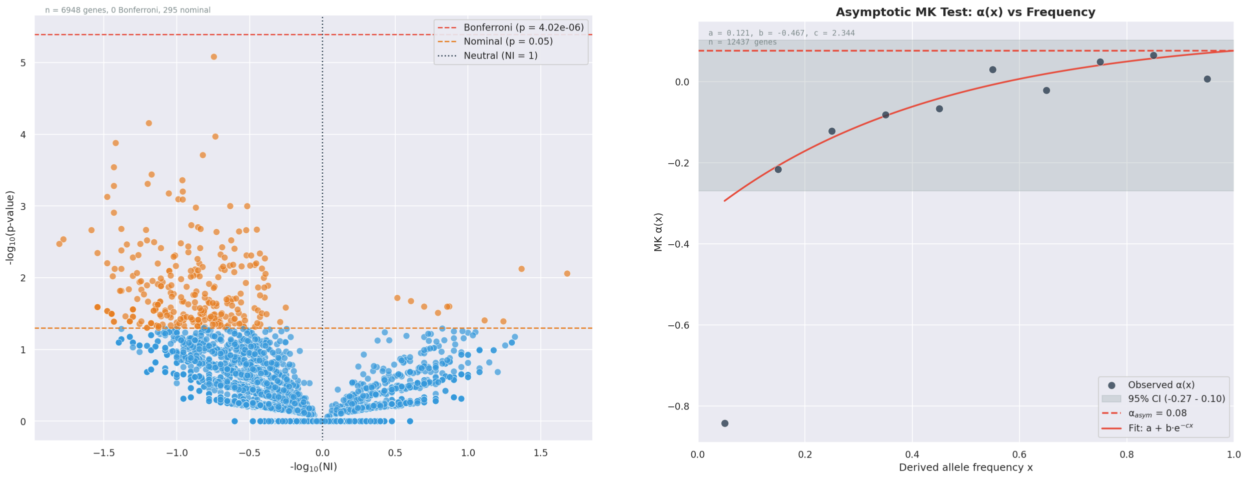


Figure S2: MK test results for 12,437 human protein-coding genes using *Pan troglodytes* as the outgroup for divergence. Left: Volcano plot from per-gene standard MK tests. Red points are significant after Bonferroni correction, while orange points are nominally significant ($p < 0.05$). The dashed lines indicate Bonferroni and nominal thresholds. Right: Asymptotic $\alpha$ curve for pooled data. Points represent the observed $\alpha$ at each derived allele frequency cutoff; the solid line represents the fitted curve. The shaded area indicates the 95% confidence interval around the fitted asymptotic $\alpha$ value.

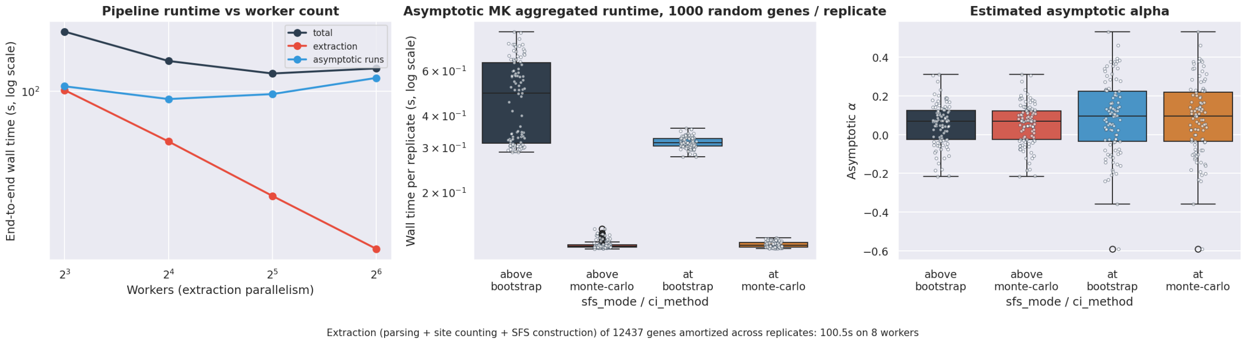


Figure S3: Aggregated-asymptotic runtime benchmark. The panels show wall time (left, log scale) and asymptotic $\alpha$ (right) across the four-cell grid of sfs-mode $\in$ {at, above} $\times$ ci-method $\in$ {monte-carlo, bootstrap}, over 100 replicates of 1,000 randomly sampled human protein-coding genes per replicate. The leftmost panel, reports end-to-end pipeline wall time at worker counts of 8, 16, 32, and 64. Per-replicate wall time covers the aggregated asymptotic-MK call only (aggregation, curve fit, and CI computation); per-gene polymorphism extraction (parsing, codon-aligning ingroup vs. outgroup, classifying nonsynonymous and synonymous polymorphism, and Nei–Gojobori site counting) runs once across all genes, is cached to disk, and is amortized across replicates—its one-time wall time is reported in the figure title. The benchmark is reproducible via the aggregated-asymptotic-runtime script in the `examples/benchmarks/` directory of the repository.
